# Bioconjugation of AuNPs with HPV 16/18 E6 antibody through physical adsorption technique

**DOI:** 10.1101/2020.01.21.899930

**Authors:** Lucy Muthoni Mwai, Mutinda C. Kyama, Caroline W. Ngugi, Edwin Walong

## Abstract

Gold nanoparticle (AuNP) bioconjugates are increasingly being utilised in biomedicine due to their low toxicity on biological tissues and unique electronic and chemical properties. They have been utilised in several biological applications namely manufacture of nanomaterials, biosensing, electron microscopy and drug delivery systems. Particularly, immuno-assays often employ gold nanoparticles (AuNPs) to enhance detection of a biological component. This paper presents a study on the bioconjugation of AuNPs with Horse Radish Peroxidase conjugated Human Papilloma Virus 16/18 Early 6 antibodies (CIP5) against Early 6 (E6) oncoprotein that is overexpressed in cervical carcinoma progression through physical adsorption. This bioconjugate can be employed in diagnostic immunoassay for cervical cancer screening. The study also demonstrated that the antibody pI, gold colloidal solution pH and amount of antibody determine the generation of stable Antibody–AuNPs bioconjugates.

## Introduction

HPV 16/18 E6 oncoprotein has been evaluated as a useful biomarker with prognostic abilities as it can detect pre-cancer and cancerous states of cervical cancer progression (1–3). A positive E6 assay, indicates high correlation to the cervical cancerous phenotype, not the potential for cervical cancer, thus high specificity in triaging patients during screening (4–6). E6 levels of expression associate directly with severity of Cervical intraepithelial neoplasia (CIN) lesions and risk of progression to invasive carcinoma (7).

Although, the current screening interventions utilizing the E6 oncoprotein from high risk HPV strains have significantly high specificity, the sensitivity is relatively low. OncoE6 test showed 69.6% sensitivity for High Grade Squamous Intraepithelial Lesion (HSIL) (CIN3+) precancerous lesions (3). HPV16/18/45 E6 test produced sensitivity and specificity of CIN2+ as 42.8% and 94.3% respectively and for CIN3+ as 54.2% and 93.8% respectively (8,9). More advanced methods of detecting E6 oncoprotein are therefore required to produce a sensitivity between 76.3 %-97.2% which is indicative of diagnostic test quality (10).

Nanobiotechnology is being used in cancer diagnostics to enhance detection. Nanotechnology refers to the utilization of materials at nanoscale ranging from 1 to 100nm in scientific fields such as chemistry, biology, physics, and engineering. At nanoscale, these materials exhibit different properties from bulk material (11). Nanobiotechnology is hence the application of nanosized tools and systems in the study of biological phenomena. It is applied in prevention (nanovaccines), diagnosis (in vivo diagnostics and in vitro diagnostics) and in treatment (drug delivery) of cancer nanooncology of human diseases (12). The types of nanomaterials include metal based (gold (Au) and silver (Ag) nanoparticles), oxides (super paramagnetic iron oxide (SPIO) nanoparticles, silver (Ag) oxide nanoparticles), liposomes, dendrimers, quantum dots, nanosphere, carbon nanotube, nanofibers, nanolayers like graphene sheets, biopolymers among others (13).

Both in vivo diagnostics and in vitro diagnostics utilize biosensors for cancer diagnosis. Biosensor devices contain a biological recognition element capable of detecting presence of a specified biological analyte such as proteins, isoenzymes, nucleic acids, metabolites or hormones and a transducer that converts the biochemical signal into quantifiable electrical signal (14). The transducers depend on the signal from biological recognition element: optical transducer (luminescence, fluorescence, interferometry and colorimetric), magnetic (electrical magnetic force), electrochemical (amperometry, potentiometry and conductometry), calorimetric (heat thermistors), mechanical (force), piezoelectric (pressure or mass changes) (14). In vivo sensors consist quantum dots, nanoshells among others to detect biochemical changes in biological systems through imaging, while in vitro sensors utilize nanoparticles, nanowires, graphene sheets among others (15).

Biosensors are increasingly being used in cancer diagnostics, primarily because in cancer development, cancer-specific biomarkers are elevated or reduced, hence their detection and identification enhances diagnosis (14). Biosensors are also applicable at point of care testing to enable screening. They exhibit high sensitivity, specificity, reproducibility and cheap instrumentation (11). Early cervical cancer diagnostics have been improved by application of nanotechnologies (16). For example; the ultra-bright fluorescent mesoporous silica nanoparticle has been utilized to detect folate (Fa) receptors on cervical cells surface and fluoresce. The brighter the fluorescence, the more the more advanced the cervical carcinoma (17). Secondly, adding monodispersed inorganic silica nanoparticles into organic dyes used to detect HR-HPV DNA using DNA microarrays has amplified DNA microarray technology signal (18). Thirdly, nanotechnology has been used to detect proteins for example the piezoelectric (contains quartz crystals) immuno-sensor that detects protein p16^INK4a^ for early cervical cancer detection (19).

This study developed a bioconjugate of gold nanoparticles with HPV 16 E6/18 E6-HRP (C1P5) antibody that can be utilised to sensitively detect E6 oncoprotein biomarker in cervical cancer screening.

## Materials and methods

### Bioconjugation preparation

The pH of 500 μl mouse IgG1 HPV 16 E6/18 E6-HRP (CP15) monoclonal antibodies (Santa Cruz Biotechnologies, USA Cat# sc-460) was determined using a pH Meter. The pH of 1000 μl 20nm colloidal spherical monodispersed citrate stabilized gold nanoparticles (Sigma Aldrich) was adjusted to a pH unit of 0.5 higher than the iso-electric point (IEP) of the antibody molecule to be adsorbed using 20 mM borate buffer pH 8.7. Using Jenway UV-VIS Spectrophotometer Model 6800, the lambda maximum of the colloidal gold was measured before bioconjugation.

### Bioconjugation optimization

A bioconjugation optimization protocol was carried out to determine the amount of antibody needed for maximum bioconjugation. This was accomplished by diluting 200 μg/ml HPV 16 E6/18 E6-HRP (CP15) antibodies in 1X PBS buffer pH 7.4 (Beijing Solarbio Science & Technology Co. Ltd) a range of 0mg/ml to 0.012mg/ml. To each dilution 100 μl of the gold solution was added. This mixture was incubated for 2 h to allow for passive adsorption (20). The absorption spectra of each mixture was obtained to confirm the concentration at which bioconjugation took place using Jenway Model 6800 UV-VIS Spectrophotometer (21).

### Bioconjugation confirmation using Spectrophotometer

The absorbance spectrum was obtained for all dilutions through a scan within the standard range of wavelengths 400-900nm. From the collected absorbance spectrum, the absorbance maximum (λ_max_) was determined.

### Preparation of HPV 16 E6/18 E6-HRP (CP15)-AuNPS conjugate for downstream processes

About 100 μl of AuNPs was added into an Eppendorf tube, the pH was adjusted to pH 7 using 1X PBS at pH 7.4. 200 μg/ml of antibody was diluted to 4μg/ml by adding 20 μl of stock to 980 μl 1X PBS buffer to make 4 μg/ml concentration of the antibody. 900 μl of the antibody concentration obtained above, was added to the 100 μl AuNPs to make a final solution of 1000 μl. This was incubated for 2 h at 4°C to allow passive adsorption. The solution was centrifuged at 2500 rpm for 1 minute to remove unbound antibody and supernatant discarded. The pelleted HPV 16 E6/18 E6-HRP (CP15) - AuNPS conjugate is resuspended in 1X PBS buffer pH 7.4. The centrifugation was repeated and the pelleted conjugate stored undiluted at 4°C.

## Results

The pH of 500μl HPV 16 E6/18 E6-HRP (CP15) antibody was obtained as pH 7.05. The pH of 1000 μl of colloidal gold solution was adjusted to pH 7.83 a pH unit of about 0.5 higher than the iso-electric point (IEP) of the antibody molecule using 20 mM borate buffer pH 8.7. The maximum absorption for the 20 nm, OD 1 gold nanoparticles with a concentration of 6.54 x 10^11^ particles/mL, a peak SPR wavelength of 518-524 nm and extinction coefficients of 9.21 x 10^8^ (M^-1^ cm^-1^ before conjugation was at 523nm as shown on Fig 1.

**Fig 1.**
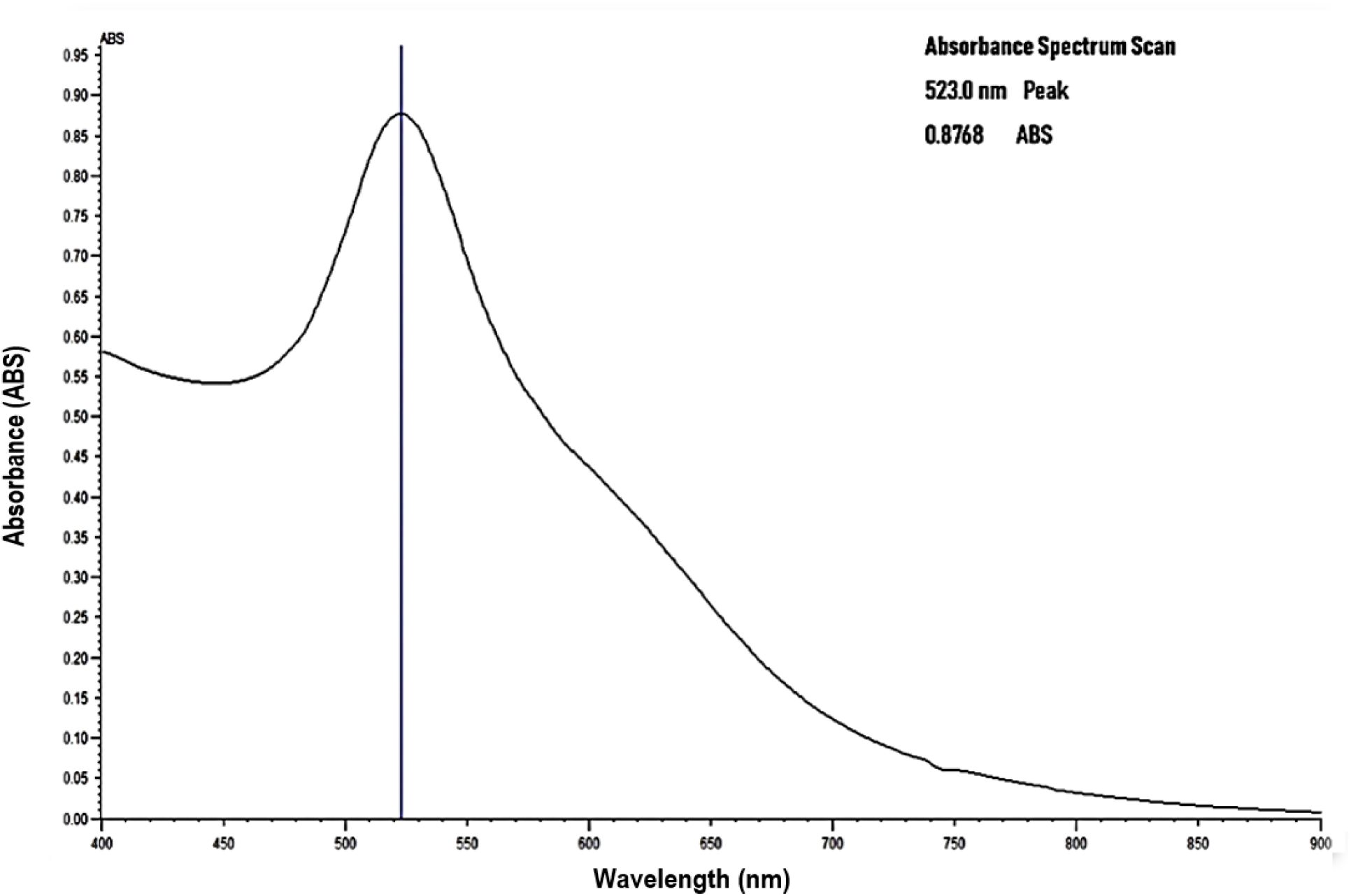
Absorption Spectra of Gold Nanoparticles before conjugation using Jenway Model 6800 UV-VIS Spectrophotometer.

After 2 h incubation, the Lambda max of 0mg/ml to 0.012mg/ml range of antibody solutions were obtained as outlined in Table 1.

**Table 1.** Showing Absorption Spectra of AuNPs after 2 h incubation of different concentrations of HPV 16 E6/18 E6-HRP mAbs with 100μl of AuNPs using Jenway Model 6800 UV-VIS Spectrophotometer.

| **HPV 16 E6/18 E6-HRP antibody concentration (µg/ml)** | **Absorption Maximum after 2 h incubation** |
| --- | --- |
| **0** | 523.0 |
| **0.3** | 519.5 |
| **0.6** | 520.0 |
| **0.9** | 520.5 |
| **1.2** | 521.0 |
| **2.0** | 522.0 |
| **4.0** | 523.5 |
| **6.0** | 522.0 |
| **8.0** | 522.0 |
| **10.0** | 521.0 |
| **12.0** | 521.5 |

From these absorption spectra, a red shift of lambda maximum by 0.5nm was observed at antibody concentration 4.0μg/ml from the initial 523.0 nm. This is further illustrated in Fig 2.

**Fig 2.**
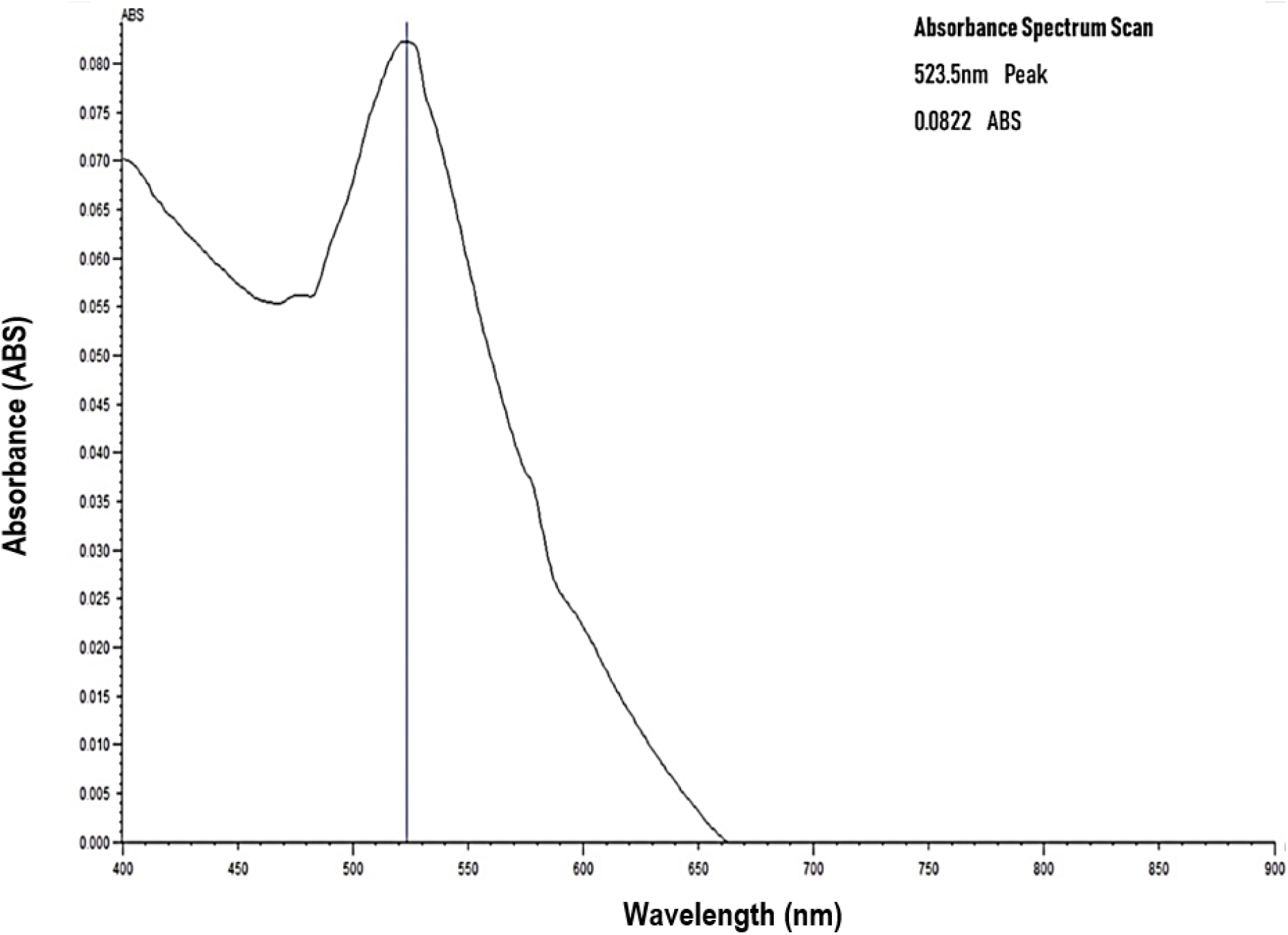
Absorption Spectra of Gold Nanoparticles after conjugation with 4.0μg/ml concentration of antibody using Jenway Model 6800 UV-VIS Spectrophotometer

## Discussion

Bioconjugation occurs upon successful binding of the monoclonal antibodies to the AuNPs surface, causing the localized surface plasmon resonance (LSPR) spectrum to red-shift that is, increase in wavelength by a few nanometers (22). The red shift indicated in this study was 0.5nm. This is attributed to the fact that the antibody replaces the negative charge citrate capping on the surface of citrate stabilized colloidal gold nanoparticles. This involved both electrostatic and hydrophobic interactions involved in passive adsorption as in Fig 3 (20).

**Fig 3.**
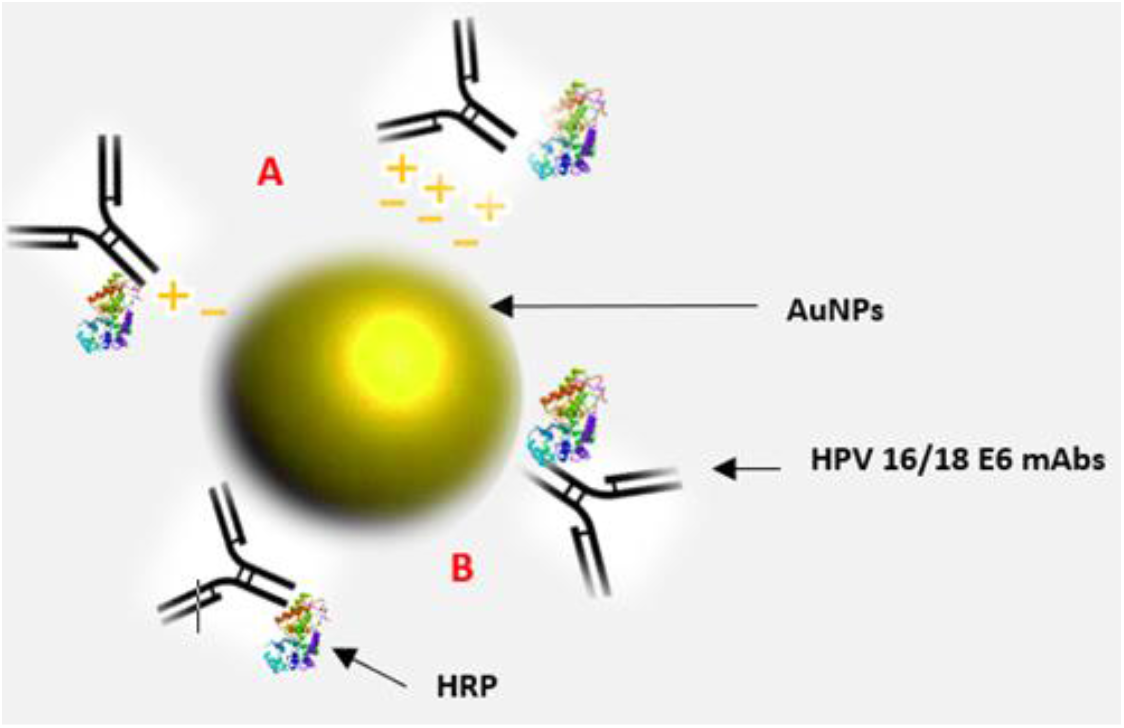
Showing bioconjugate: HPV 16 E6/18 E6-HRP (CP15)-AuNPs. It comprises 20nm citrate stabilized monodispersed spherical AuNPs passively adsorbed to horseradish peroxidase (HRP) enzyme labelled HPV 16/18 E6 (C1P5) mouse monoclonal antibodies through (A) electrostatic and (B) hydrophobic interactions

To ensure optimal bioconjugation, there were critical steps involved. Firstly, there was adjustment of the pH of colloidal gold solution to pH 7.83, a pH unit of about 0.5 higher than the iso-electric point (IEP) of the antibody molecule to be adsorbed whose isoelectric point is about pH 7.00 (23). This is critical because hydrophobic interactions are maximal at IEP of antibody hence maximal bioconjugation was achieved (24,25).

Secondly, the concentration of antibody needed was optimized through titrations of the antibody as too little antibody adsorbed to the gold surface, will cause aggregation upon addition of electrolytes present in standard buffers. Bioconjugation occurred at 4.0μg/ml HPV 16 E6/18 E6-HRP (CP15) antibody concentration. The bioconjugate formed was prepared and pelleted conjugate stored undiluted at 4°C for use in E6 oncoprotein biomarker detection in cervical cancer screening. It can be stored for 12-18 months.The absorption spectrum obtained after bioconjugation was fairly smooth (Figure 2) due to possible antibody contamination with small contaminants such as sodium azide. Dialysing the antibody with a buffer solution is highly recommended in subsequent analysis. It could also arise from contamination of gold colloidal solution during pipetting hence aliquoting the gold solution during use is to be maintained. Despite the challenges, passive adsorption of gold nanoparticles to antibody was achieved, and testing the antibody sensitivity on E6 protein will further verify the findings.

Early cervical cancer diagnostics have been improved by application of nanotechnologies (16). Particularly, immuno-assays often employ gold nanoparticles (AuNPs) to enhance detection of a biological component. Gold nanoparticle (AuNP) bioconjugates are increasingly being utilised in biomedicine due to their low toxicity on biological tissues and unique electronic and chemical properties such as unique surface characteristics, optical properties and consistency (26). The Bioconjugate herein, uses an antibody conjugated to HRP thus conjugation to AuNPs will further enhance visual enzyme linked immunosorbent assay (ELISA) based colorimetric detection of the E6 oncoprotein analyte (27).

## Conclusion

It was possible to conjugate mAbs against HPV 16/18 E6 oncoprotein to 20nm citrate stabilized gold nanoparticles through passive adsorption. The bioconjugate formed was HPV 16 E6/18 E6-HRP (CP15)-AuNPs. Currently, there is no report on the development a gold nanoparticle bioconjugate with mAbs against the E6 protein of HPV 16/18 to enhance the signal of the immunoassay involving the E6 protein and the antibody thereof. This HPV 16 E6/18 E6-HRP (CP15)-AuNPs conjugate can be utilized in immunoassays for further verification of the bioconjugate’s bio-functionality.

## Acknowledgements

We would like to express our gratitude to the staff at Molecular Biology and biotechnology PAUSTI Laboratory and University of Nairobi Histopathology laboratory for their support and creating a conducive environment in which the lab investigations were carried out, Dr. Dickson Andala for the support in the preliminary stages of the research study and James G. Maina for valuable feedback during the research period.

